## Supplemental materials for "Climate and urbanization drive changes in the habitat suitability of *Schistosoma mansoni* competent snails in Brazil"

**Occurrence points - collection methods**

Snails were collected using the standard technique recommended by the Brazilian Ministry of Health Schistosomiasis Surveillance and Control Program (2008). The collection points were georeferenced with a Garmin ETrex Summit® GPS. The three Neotropical hosts – *Biomphalaria glabrata*, *B. straminea,* and *B. tenagophila* – were identified simultaneously by morphology and DNA barcoding according to[1–3].

**Environmental data processing**

We used the CHELSA dataset to calculate 10-year climatologies spanning four decades (1980-1989, 1990-1999, 2000-2009, 2010-2017)[4]. We calculated these variables using the monthly precipitation and temperature from CHELSA dataset version 2.1 and aggregated it to get 10-year climatologies.

Globally, schistosomiasis is considered a rural disease but has been observed close to or in urban areas in Brazil. To measure the impact of urbanization on *Biomphalaria* occurrence, and because a urban:rural dichotomy does not capture the complexity of urbanization in Brazil[5], we derived a rural to urban gradient, where 0 indicated that the snail was observed in an urban area and 1 indicated the snail was in a rural area, and continuous values from 0-1 indicate the relative location of the snail along this gradient. We defined urban and rural areas using population density and population size as a proxy. As population-based definitions of urban are inconsistent, we defined urban areas based on two different definitions and conducted separate analyses for each one. First, we defined an urban area based on an aggregation of urban definitions throughout South America: an urban area had population densities from 300-1500 people within 1 $km^2$ with contiguous pixels totalling at least 2500 people. We then defined urban pixels based on the United Nations definition, which uses the same pixel density but with contiguous pixels totalling at least 5000 people[6]. Finally, we included high density urban areas in both analyses: > 1500 people per grid cell with contiguous grid cells totalling more than 150,000 people[6]. We used WorldPop to estimate population size and cumulative cost mapping – where every pixel is assigned the total cost of the lowest cost path (distance) to an urban pixel – to estimate the distance to the urban pixel for points collected between 2000 - 2020[7]. WorldPop data is not available before 2000; as such, we derived population estimates from 1990 - 1999 by linearly interpolating between the 2000 WorldPop raster and the Global Human Development Layer, Population Grid 1975 raster[8]. We used linear interpolation opposed to exponential, which is the trend observed in long-term change of population size, as the study is over a relatively short period of time and a linear trend is adequate to describe the population trend. To calculate the final variable - location along the rural to urban gradient - we divided distance to an urban area by distance to a rural area and standardized it using a max-min transformation. We present results based on the definition of urbanization based on an aggregation of South America definitions, but include results based on the UN's definition. Model results and interpretation are comparable across models (supplement pp 7, 17-22).

To estimate land use/land cover, we used the MAPBIOMAS land-cover rasters collection 7.0 version 2 (annual at 30m resolution) to calculate the percent cover of temporary crops and mosaic of use (including urban farms), two land-use types known to influence snail presence, within the 1$km^2$ grid cell during the year the data point was collected[9]. Temporary crops did not include temporary crops associated with monoculture (soybeans, sugar cane, rice, cotton), but did include other crops with a vegetative cycle of less than one year, this includes lettuce, which has been found to create snail habitat in urban farms (Tuan, personal communication). Mosaics of use include areas where it is not possible to distinguish agricultural land from pasture and, in urban areas, includes cultivated vegetation and natural forest and non-forest vegetation[9].

To estimate important hydrologic and typologic covariates, we used the Global Surface Water dataset (30m resolution) to estimate the water occurrence (the frequency with which water occurred), mean water recurrence (the frequency with which water returns year to year), mean water seasonality (number of months water is present)[10]. We then used the MERIT Hydro data (100m resolution) to calculate the mean upscale drainage area (upa) and maximum height above the nearest drainage (hnd)[11]. The distance to the nearest river was estimated using the WWF HydroSHEDS Free Flowing River dataset and cumulative cost mapping[12]. We described the topography (mean elevation, mean slope, and mean aspect) using the NASA NASADEM Digital Elevation dataset (30m resolution)[13]. Finally, we used the OpenLandMap dataset (250 m resolution) to estimate mean soil clay[14], sand[15], water[16], and carbon content[17], pH[18], and soil bulk density[19] at a depth of 0mm. Hydrology, topography, and soil property datasets were static or available at aggregations across the study period. All environmental data was processed and aggregated to 1km$^2$ grid cells around each data point using Google Earth Engine[20] and the GEE Python API in Google Co-Laboratory, with the exception of CHELSA, which was downloaded from their AWS repository[21,22].

Highly correlated features were removed to enhance model interpretation using the “findCorrelation” function in the R package *caret*[23]*.* We removed the feature if there was > 0.75 correlation with another feature. Between the two features, we removed the feature that reduced the highest number of collinear features in the model (e.g., if soil clay content was correlated with soil bulk density, but soil bulk density was correlated with both soil pH and soil sand content, we removed soil bulk density). Variables retained in the final analysis are outlined in (Appendices 3-5; 12-15). We removed the feature if there was >0.75 correlation with another feature. Our final variables were: mean diurnal range of temperatures averaged over 1 year (diurnal temp range), ratio of diurnal variation to annual variation in temperatures (isothermality), mean daily air temperatures of the wettest quarter (temp wettest qt), annual precipitation amount (annual precip), precipitation amount of the wettest month (precip wettest month), precipitation amount of the driest month (precip driest month), coefficient of variation of monthly precipitation (precip seasonality), mean monthly precipitation amount of the wettest quarter (precip wettest qt), distance to the nearest pixel with human population density >300 per 1km$^2$ (distance to urban pixel), distance to the nearest pixel with human population density >1500 per 1km$^2$ (distance to high density urban pixel), percent coverage of temporary crops, percent cover of agricultural mosaic, mean water recurrence, mean upstream drainage area (upa), mean height above nearest drainage (hnd), mean aspect, soil clay content, soil sand content, soil water content, soil carbon content, soil pH, soil bulk density, and distance to nearest river.

**Extreme Gradient Boosted Regression Tree SDMs - Spatial CV**

Spatial cross-validation (5-fold) was used to evaluate model performance to estimate the model's predictive power in geographic regions not used for training so as to avoid inflating model performance values due to spatial autocorrelation of environmental predictors. First, occurrence points were split into folds based on their geographic coordinates using the R package *blockCV*[24]. For each iteration (one per fold), one fold was held out as the test set, and the model was trained on the remaining data. This procedure was repeated five times for each fold. Model parameters were tuned and trained using Bayesian optimization via the package *bayesianOptimization*[25]. Specifically we tuned: the learning parameter (eta; controls how much information from a new tree will be used for boosting), maximum depth of the trees, minimum number of observations in a terminal node, the fraction of data used to grow each tree, and the fraction of features used to train each tree for each bootstrapping iteration. We set gamma to 0 and the ratio number of positive to negative classes to 2 as there were 2x as many background samples (0) as presence samples (1). Model performance was then evaluated based on how well the model predicted data within the held-out fold. Specifically, model sensitivity (i.e., the proportion of occurrences the model correctly identifies as occurrences), model specificity (i.e, the proportion of pseudo-absences that the model correctly identifies as pseudo-absences), and model area-under-the-curve (AUC; i.e., a measure that calculates how well the model correctly identifies occurrence points versus pseudo-absence points, where AUC ≤ 0.5 indicates the model performs no better than a coin flip) were calculated using the R package *pROC*[26] and *caret*[27]. Model training and validation were conducted using data from 2000-2020. The final model performance scores were averaged across the five folds. Final steps of data cleaning and machine learning analyses were done in R v 4.0.2.

**Supplementary Table 1.** Features considered for the species distribution model. Features in bold are the ones selected for the final model. Feature selection was based on examining the pairwise correlations and removing features so that no pairwise correlation was > 0.75. For variables removed from the analysis, the variables they were highly correlated with are included in the last column. For the complete description of each variable, please refer to the data source.

| Feature | Short definition | Spatial res ($km^2$) | Temporal resolution | Source | Correlation with? |
|---|---|---|---|---|---|
| bio1 | Mean annual air temp averaged over 1 year (C) | 1 | 10-year average | CHELSA | bio8, bio9 |
| **bio2 (diurnal temp range)** | Mean diurnal range of temps averaged over 1 year (C) | 1 | 10-year average | CHELSA | |
| **bio3 (isothermality)** | Ratio of diurnal variation to annual variation in temps (C) | 1 | 10-year average | CHELSA | |
| bio4 | Standard deviation of the monthly mean temp (C) | 1 | 10-year average | CHELSA | bio3, bio9 |
| bio4a | CV of the month mean temp (C) | 1 | 10-year average | CHELSA | bio3 |
| bio5 | Mean daily maximum air temp of the warmest month (C) | 1 | 10-year average | CHELSA | bio8, bio9 |
| bio6 | Mean daily minimum air temp of the coldest month (C) | 1 | 10-year average | CHELSA | bio9 |
| bio7 | Annual range of air temp (C) | 1 | 10-year average | CHELSA | bio2 |
| **bio8 (temp wettest qt)** | Mean daily mean air temps of the wettest quarter (C) | 1 | 10-year average | CHELSA | |
| **bio9 (temp driest qt)** | Mean daily mean air temps of the driest quarter (C) | 1 | 10-year average | CHELSA | |
| bio10 | Mean daily mean air temps of the warmest quarter (C) | 1 | 10-year average | CHELSA | elevation, bio8, bio9 |
| bio11 | Mean daily mean air temps of the coldest quarter (C) | 1 | 10-year average | CHELSA | bio8, bio9 |
| **bio12 (annual precip)** | Annual precipitation amount ($kg\ m^{-2}\ year^{-1}$) | 1 | 10-year average | CHELSA | |

| | | | | | |
|---|---|---|---|---|---|
| **bio13 (precip wettest month)** | Precipitation amount the wettest month (kg $m^{-2}$ $year^{-1}$) | 1 | 10-year average | CHELSA | |
| **bio14 (precip driest month)** | Precipitation amount the driest month (kg $m^{-2}$ $year^{-1}$) | 1 | 10-year average | CHELSA | |
| **bio15 (precip seasonality)** | Coefficient of Variance of monthly precipitation (kg $m^{-2}$ $year^{-1}$) | 1 | 10-year average | CHELSA | |
| **bio16 (precip wettest qt)** | Mean monthly precipitation amount of the wettest quarter (kg $m^{-2}$ $year^{-1}$) | 1 | 10-year average | CHELSA | |
| bio17 | Mean monthly precipitation amount of the driest quarter (kg $m^{-2}$ $year^{-1}$) | 1 | 10-year average | CHELSA | bio14, bio15 |
| bio18 | Mean monthly precipitation amount of the driest quarter (kg $m^{-2}$ $year^{-1}$) | 1 | 10-year average | CHELSA | bio13 |
| bio19 | Mean monthly precipitation amount of the coldest quarter (kg $m^{-2}$ $year^{-1}$) | 1 | 10-year average | CHELSA | bio14, bio15 |
| **Distance to urban pixel** | Distance to the nearest pixel with a human population > 300, calculated using cumulative cost mapping | 1 (the original data source was 0.1) | annual | WorldPop & Global Human Settlement Layer, Population Grid | |
| **Distance to high density urban pixel** | Distance to the nearest pixel with a human population > 1500, calculated using cumulative cost mapping | 1 (the original data source is 0.1 | annual | WorldPop & Global Human Settlement Layer, Population Grid | |
| **% coverage of temporary crops** | % of pixel covered with temporary crops () | 1 (the original data source is 0.03) | annual | MAPBIOMAS | |
| **% cover of agriculture pasture mosaic** | % of pixel covered with agricultural pasture mosaic () | 1 (the original data source is 0.03) | annual | MAPBIOMAS | |
| Mean water occurrence | The frequency with which water was present | 0.03 | calculated from 1984-2021 | JRC Global Surface Water Mapping Layers | water recurrence |

| | | | | | |
|---|---|---|---|---|---|
| **Mean water recurrence** | the frequency with which water returns year to year | 0.03 | calculated from 1984-2021 | JRC Global Surface Water Mapping Layers | |
| Mean water seasonality | # of months water is present | 0.03 | calculated from 1984-2021 | JRC Global Surface Water Mapping Layers | water recurrence |
| **Mean upstream drainage area (upa)** | Surface area of flow accumulation area | 0.09 | average from 1987 - 2017 | MERIT Hydro: Global Hydrography Dataset | |
| **Height above nearest drainage (hnd)** | The vertical height of any point on the landscape from the nearest stream surface or bed | 0.09 | average from 1987 - 2017 | MERIT Hydro: Global Hydrography Dataset | |
| **Mean elevation** | | 0.03 | static | NASADEM: NASA Digital Elevation | |
| Mean slope | | 0.03 | static | NASADEM: NASA Digital Elevation | hnd |
| **Mean aspect** | | 0.03 | static | NASADEM: NASA Digital Elevation | |
| **Soil clay** | Soil clay content in % (kg/kg) at 0 cm | 0.25 | Average from 1950-2018 | OpenLandMap Clay Content | |
| **Soil sand** | Sand sand content in % (kg/kg) at 0 cm | 0.25 | Average from 1950-2018 | OpenLandMap Sand Content | |
| **Soil water** | Soil water content (volumetric %) for 33kPA at 0 cm | 0.25 | Average from 1950-2018 | OpenLandMap Soil Water Content at 33kPa | |
| **Soil carbon** | Soil organic carbon content in x 5g / kg | 0.25 | Average from 1950-2018 | OpenLandMap Soil Organic Carbon Content | |
| **Soil pH** | Soil pH in $H_2O$ at 0 cm | 0.25 | Average from 1950-2018 | OpenLandMap Soil pH in $H_2O$ | |
| **Soil bulk density** | Soil bulk density (fine earth) 10 x kg / $m^3$ at 0 cm | 0.25 | Average from 1950-2018 | OpenLandMap Soil Bulk Density | |
| **Distance to river** | Distance to the nearest river, calculated using cumulative cost mapping | 1 | Snapshot in 2000 | WWF HydroSheds Free Flowing River Networks v1 | |

**Table 1. Model performance metrics for the three *Biomphalaria* species based on the models with the UN definition of urbanization.** The results for 5-fold cross validation include the mean, with the 95% confidence interval in parentheses. For model validation (5-fold cross validation), the number of background points were 2x the number of occurrence points. For the hindcast model testin, the number of background points used for hindcasting, which were based on retaining one point per 1km$^2$ grid cell, was 1688, 1691, and 1684 for *B. glabrata*, *B. straminea*, and *B. tenagophila,* respectively. Notably, these points were not used to train the model but only to test model predictions, therefore, class imbalance does not impact results.

| Species | No. occurrence points | AUC | Sensitivity | Specificity |
|---|---|---|---|---|
| ***Model validation: 5-fold spatial cross-validation*** | | | | |
| *B. glabrata* | 165 | 0.85 (0.80 - 0.91) | 0.86 (0.75 - 0.98) | 0.77 (0.67 - 0.87) |
| *B. straminea* | 283 | 0.84 (0.76 - 0.93) | 0.79 (0.61 - 0.98) | 0.79 (0.70 - 0.88) |
| *B. tenagophila* | 173 | 0.84 (0.72 - 0.97) | 0.89 (0.80 - 0.98) | 0.77 (0.60 - 0.93) |
| ***Hindcasting: testing model on 1990-1999 data*** | | | | |
| *B. glabrata* | 40 | 0.84 | 0.85 | 0.74 |
| *B. straminea* | 33 | 0.88 | 0.97 | 0.71 |
| *B. tenagophila* | 28 | 0.82 | 0.89 | 0.64 |

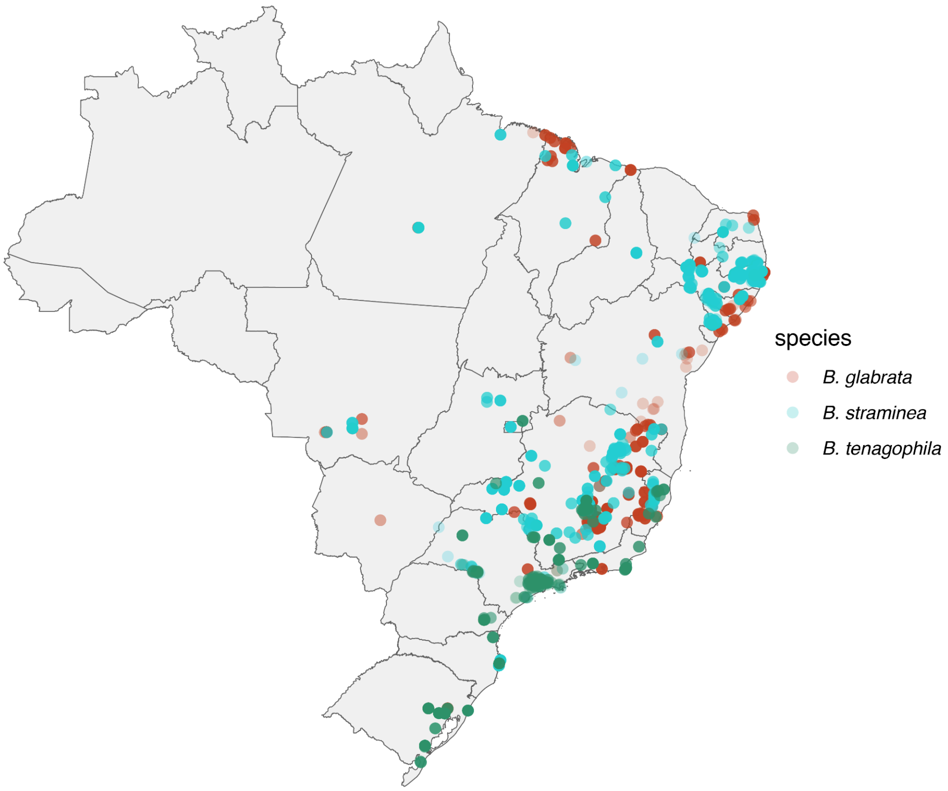


**Supplementary Figure 1. Thinned occurrence points of the three snail species.**

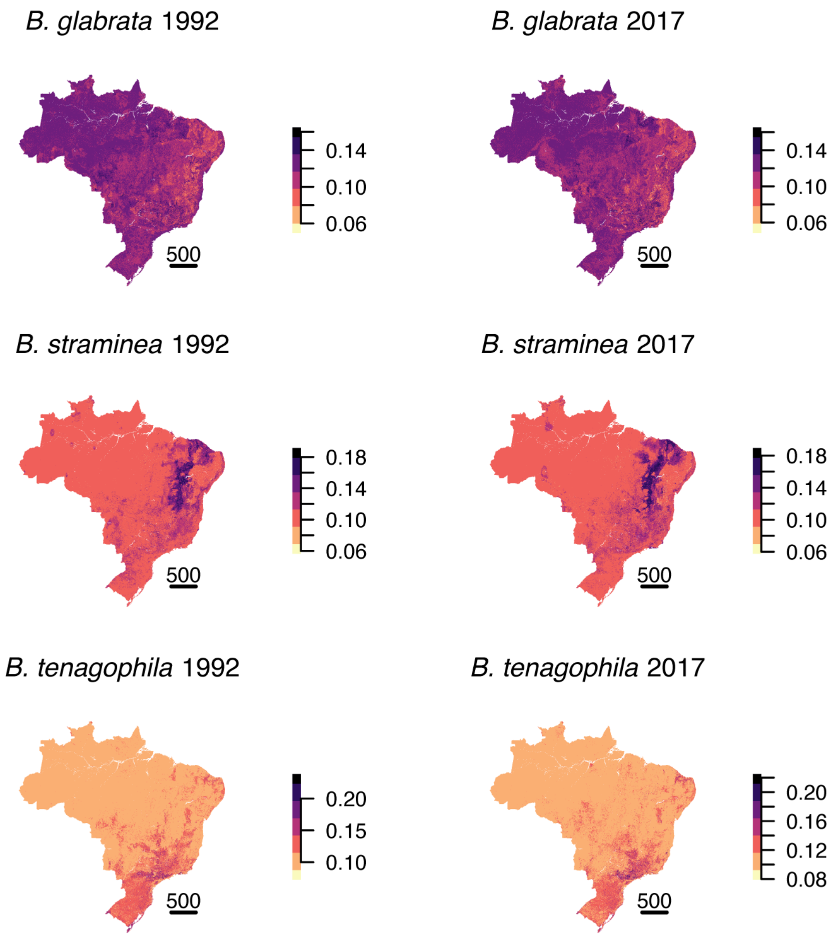


**Supplementary figure 2. Standard deviation (uncertainty) in model historical and current predicted distributions.** Standard deviation was calculated based on the probability of occurrence for 25 bootstrapping iterations (the model fit with a random, stratified subset of 80% of the data 25x).

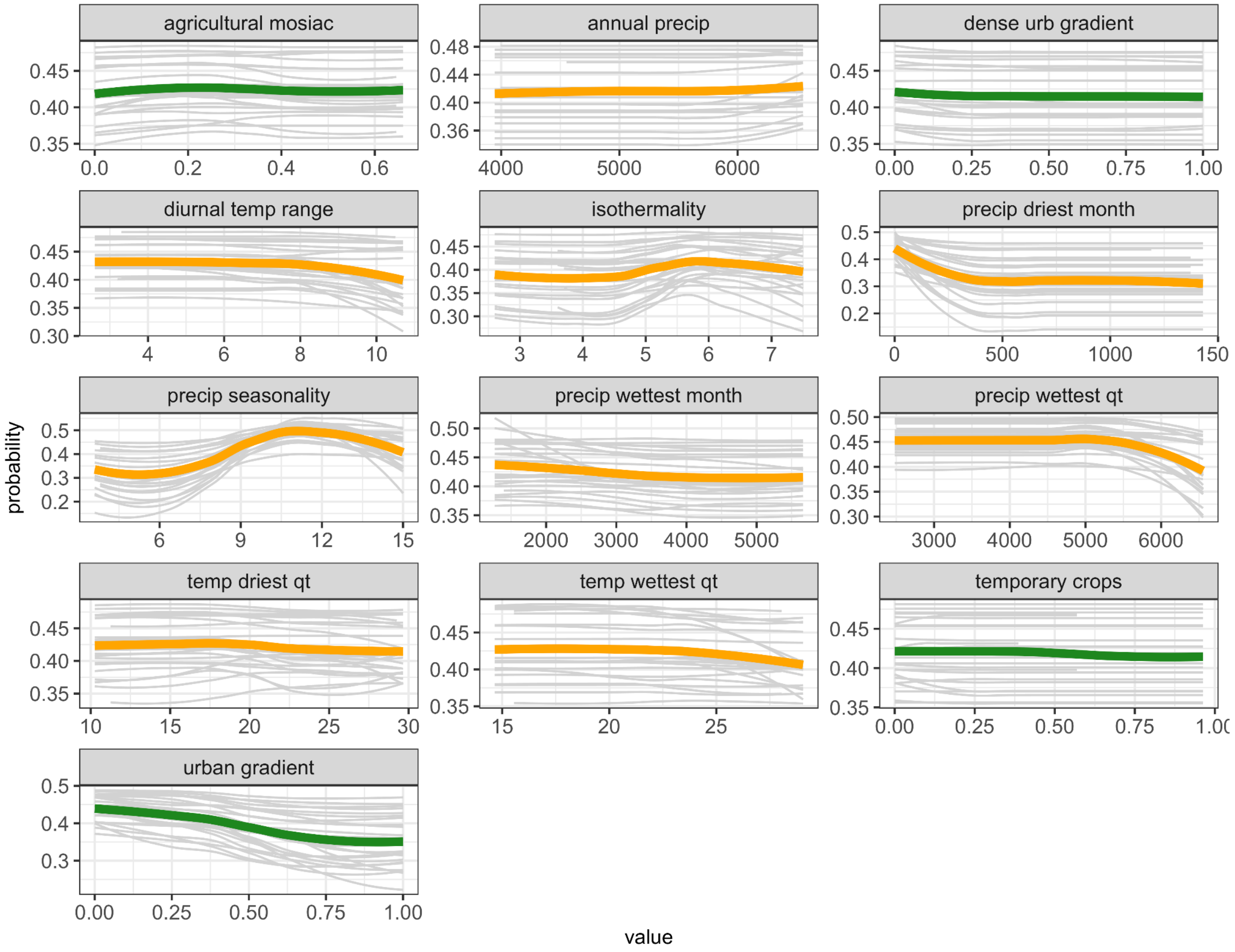


**Supplementary figure 3. *B. glabrata* partial dependence plots for climate and land-use.** Gray lines represent PDPs of each bootstrapping iteration, color lines represent average across iterations (orange = climate feature, green = land-use feature). Plots where the line looks flat indicate features with minimal contribution to feature importance.

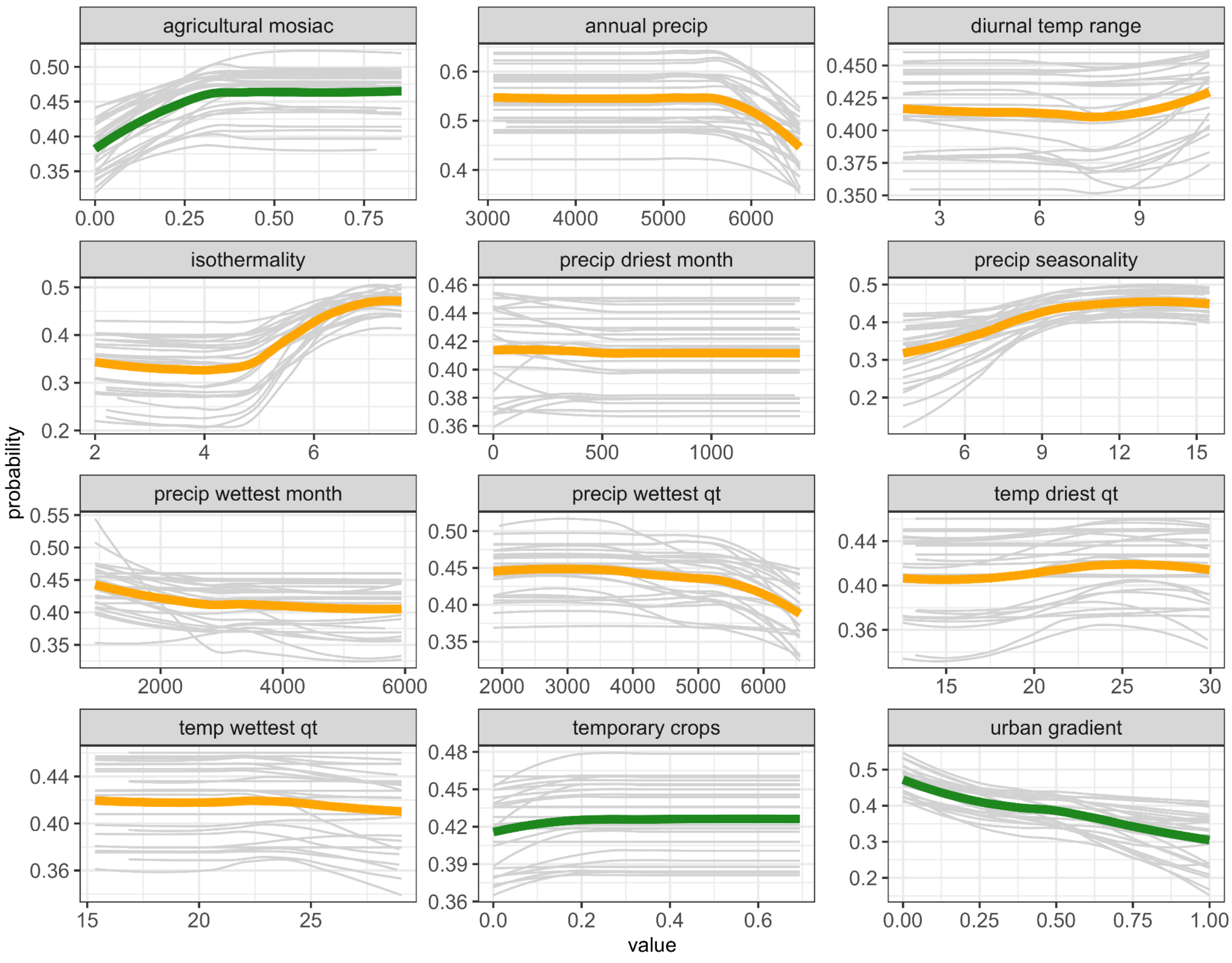


**Supplementary figure 4. *B. straminea* partial dependence plots for climate and land-use.** Gray lines represent PDPs of each bootstrapping iteration, color lines represent average across iterations (orange = climate feature, green = land-use feature). Plots where the line looks flat indicate features with minimal contribution to feature importance.

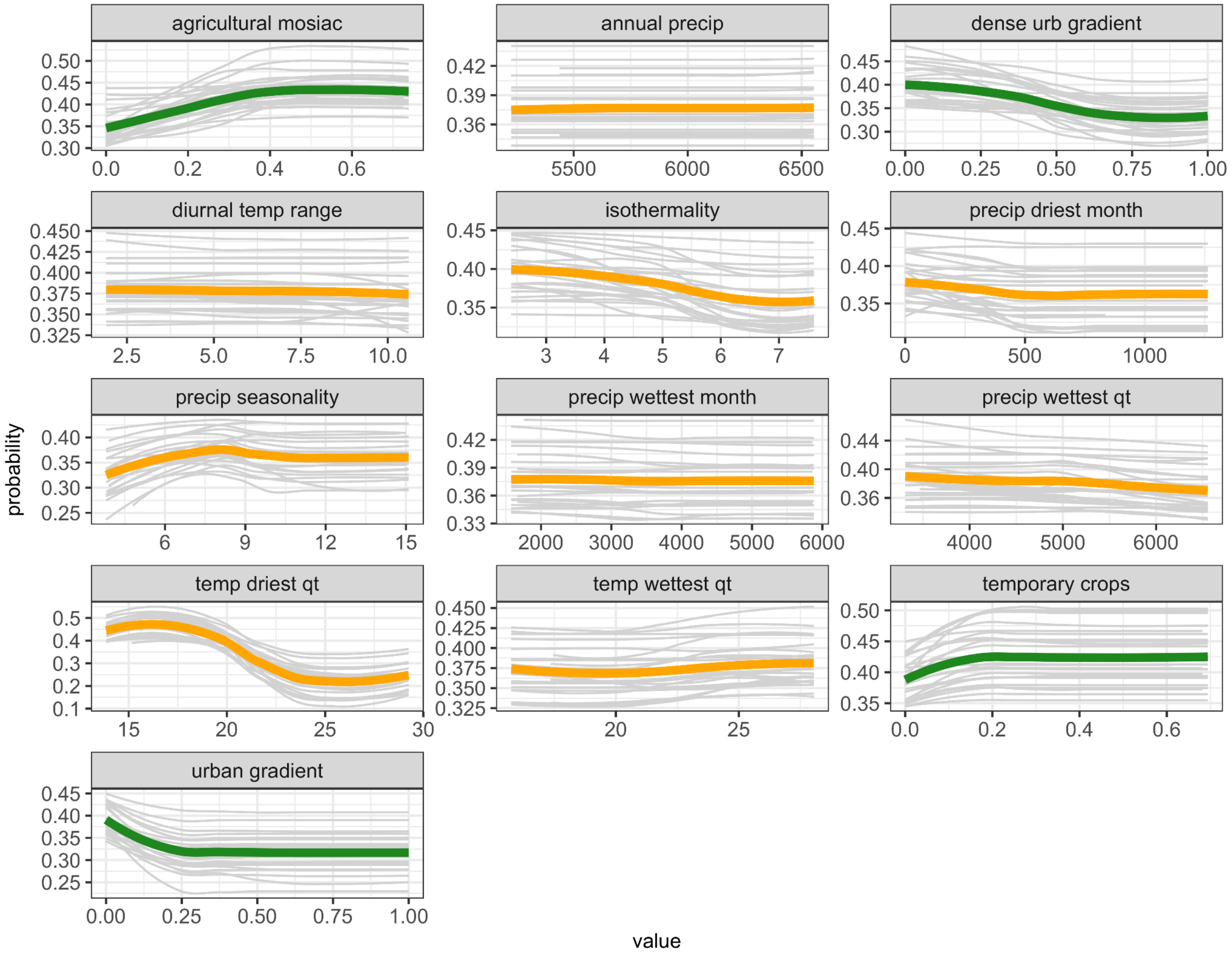


**Supplementary figure 5. *B. tenagophila* partial dependence plots for climate and land-use.** Gray lines represent PDPs of each bootstrapping iteration, color lines represent average across iterations (red = climate feature, green = land-use feature). Plots where the line looks flat indicate features with minimal contribution to feature importance.

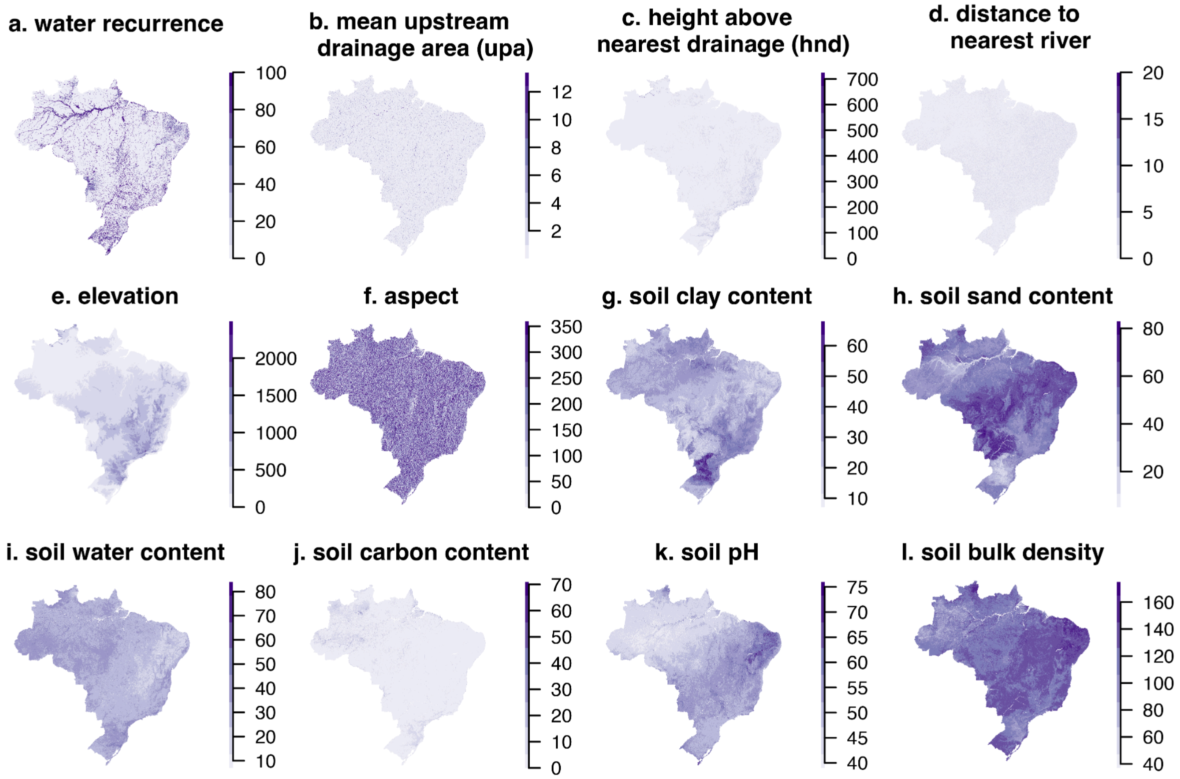


**Supplementary figure 6. Rasters for time-invariant data included in the model.**

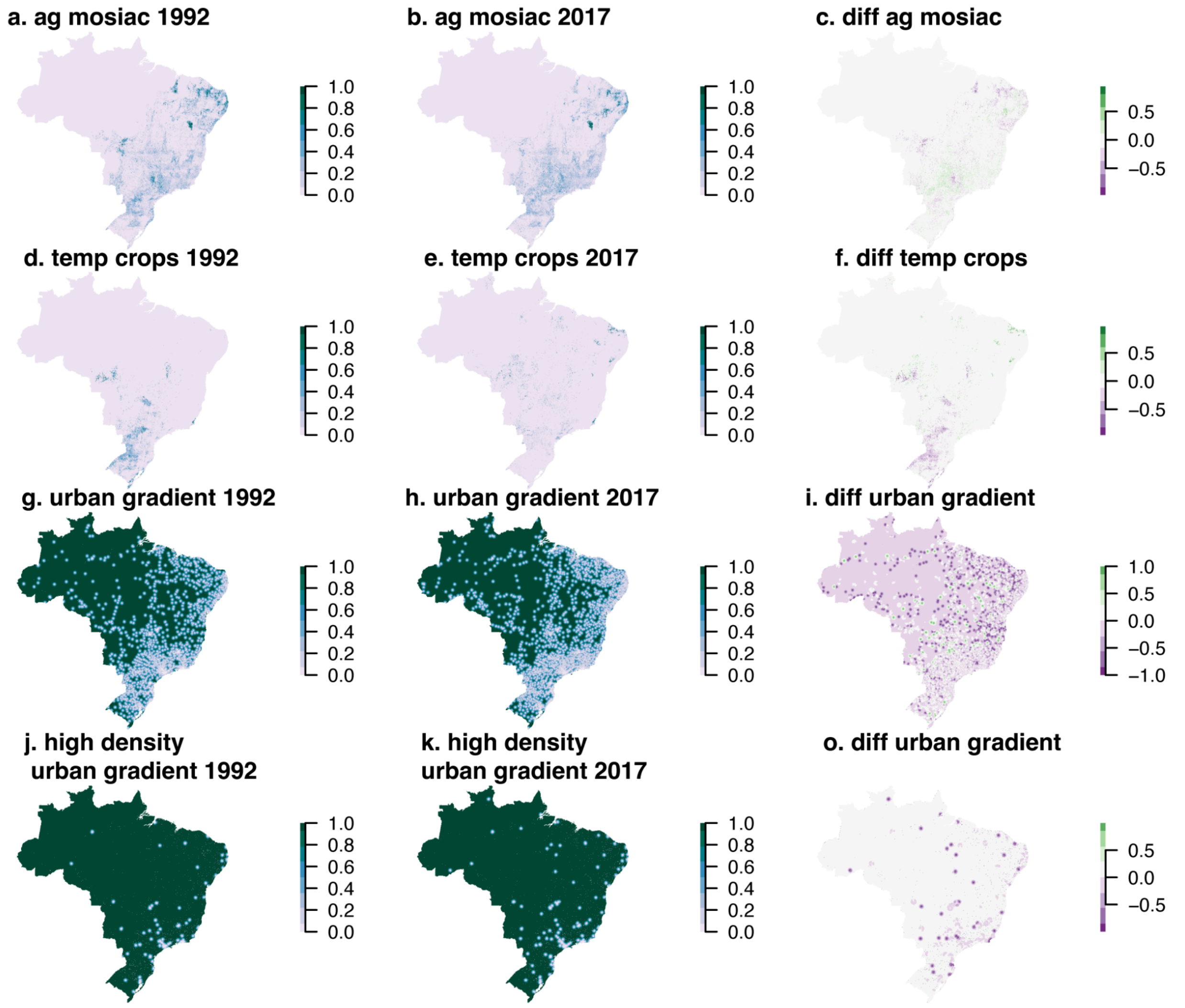


**Supplementary figure 7. Rasters for land-use change related variables included in the final model and their difference over the study period.** Land-use data varied on an annual scale but only years 1993 and 2017 are shown here (the time points used in the counterfactual analysis).

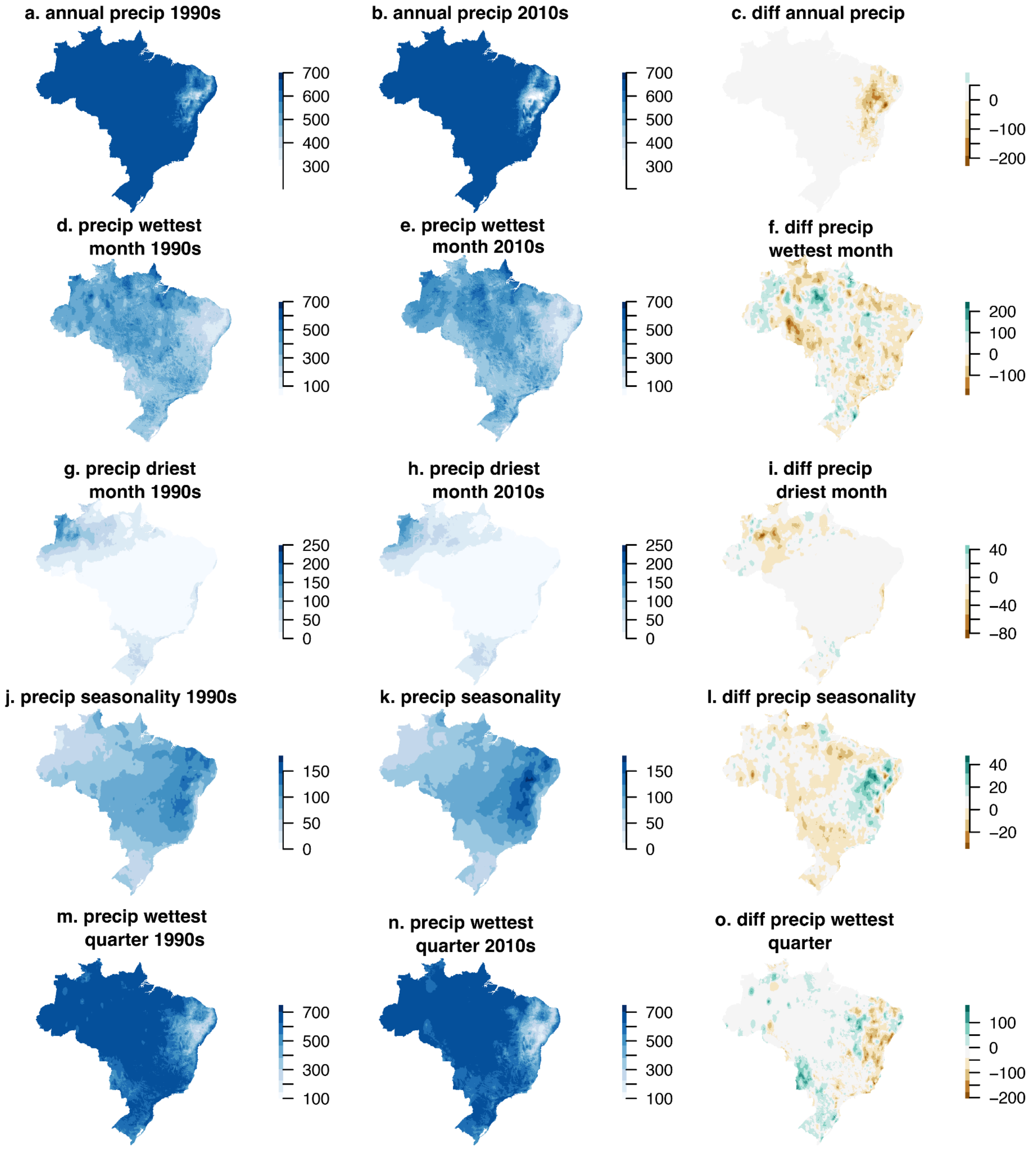


**Supplementary figure 8. Rasters for precipitation variables included in the final model and their difference over the study period.** Precipitation data varied on decadal scale but only years 1993 and 2017 are shown here (the time points used in the counterfactual analysis).

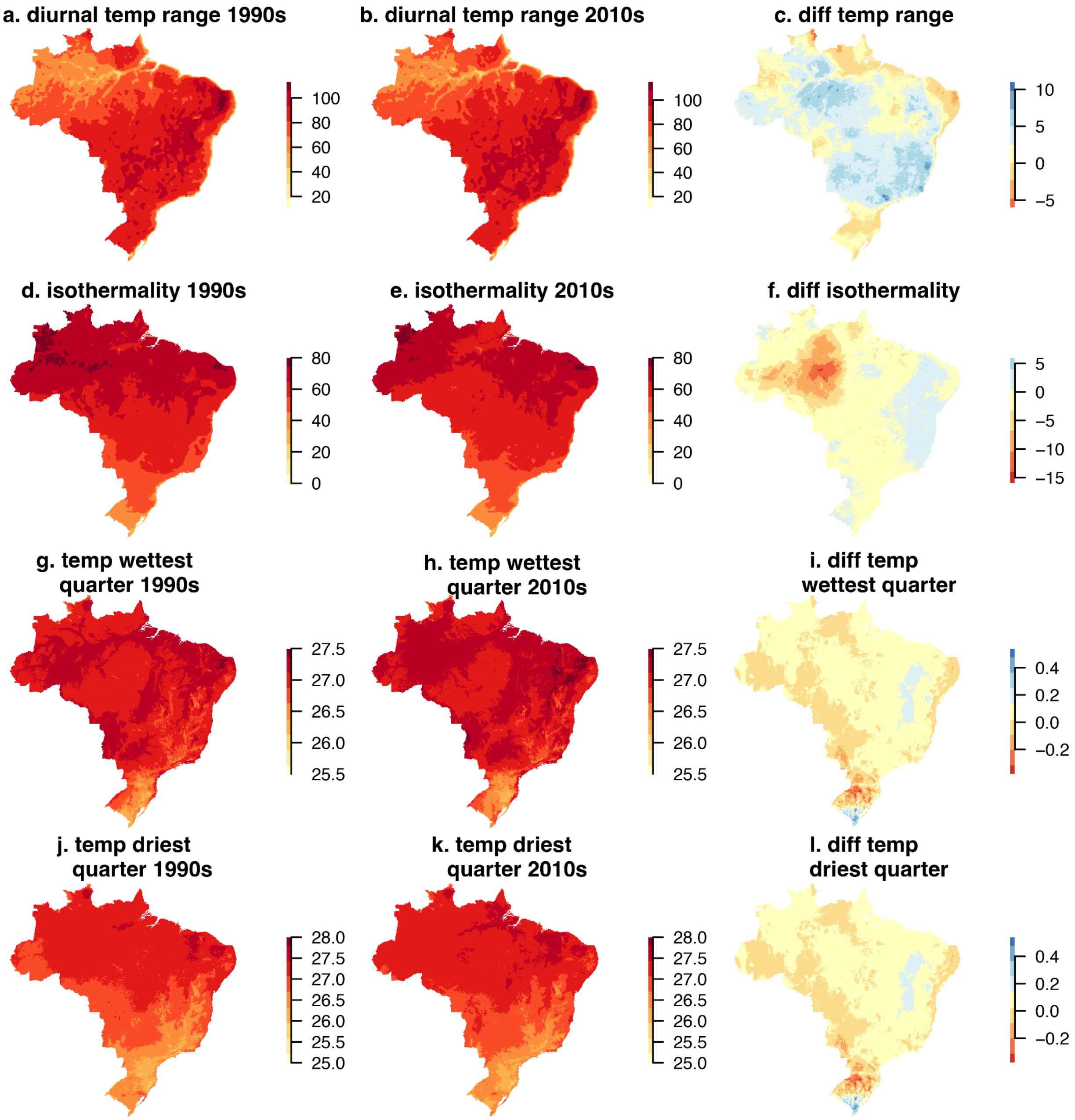


**Supplementary figure 9. Rasters for temperature variables included in the final model and their difference over the study period.** Temperature data varied on decadal scale but only years 1993 and 2017 are shown here (the time points used in the counterfactual analysis).

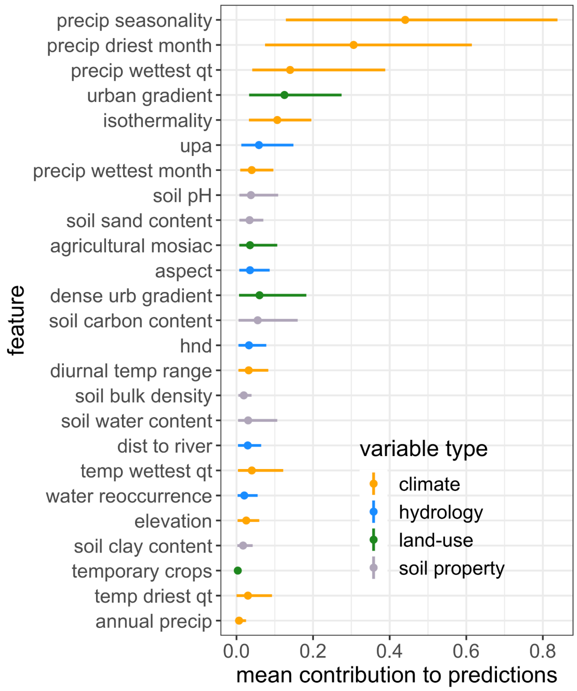


**Supplementary figure 10. Contributions to predictions for the *B. glabrata* model when defining an urban area based on the UN definition.** Points are the absolute value of the mean contribution to predictions (mean |SHAP Value|) for the covariate across all data points (i.e., global feature contribution), bars represent the 95% confidence interval.

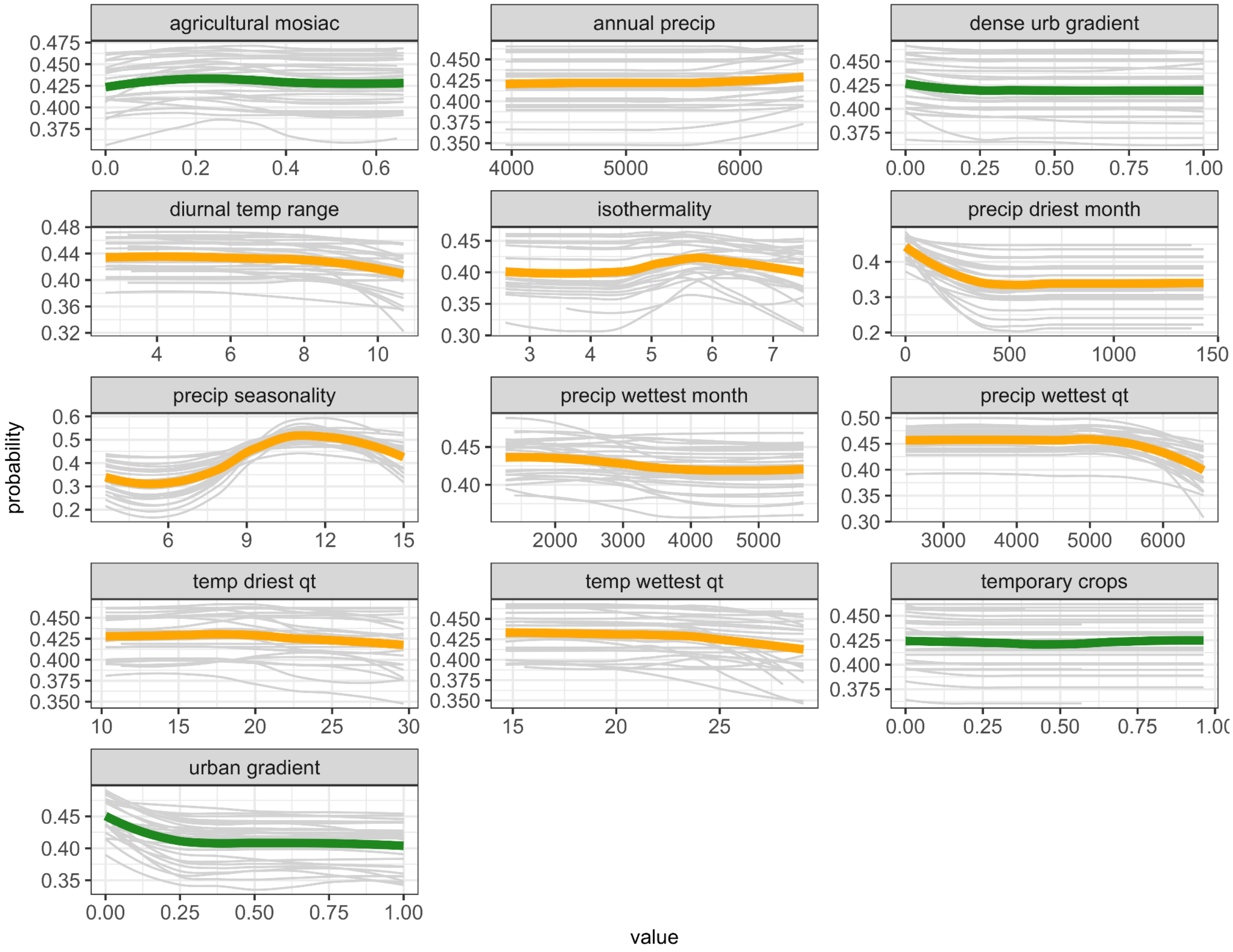


**Supplementary figure 11.** ***B. glabrata*** **partial dependence plots for climate and land-use when the urban area is defined based on the UN definition.** Gray lines represent PDPs of each bootstrapping iteration, color lines represent average across iterations (red = climate feature, green = land-use feature). Plots where the line looks flat indicate features with minimal contribution to feature importance.

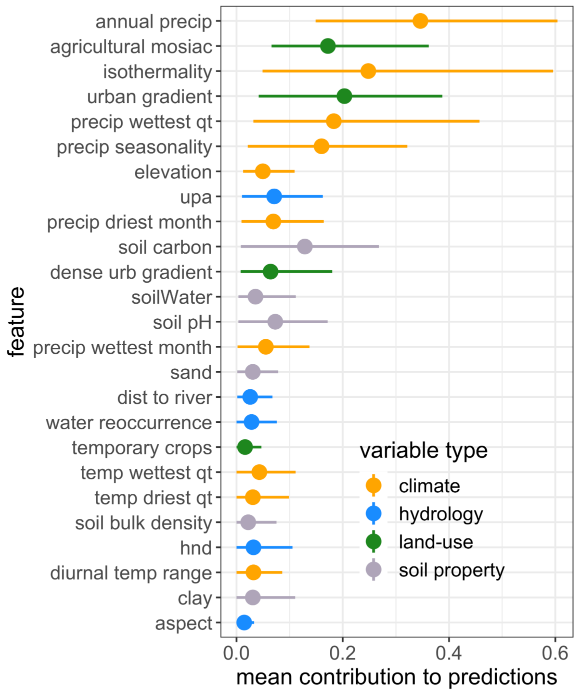


**Supplementary figure 12. Contributions to predictions for the *B. straminea* model when defining an urban area based on the UN definition.** Points are the absolute value of the mean contribution to predictions (mean |SHAP Value|) for the covariate across all data points (i.e., global feature contribution), bars represent the 95% confidence interval.

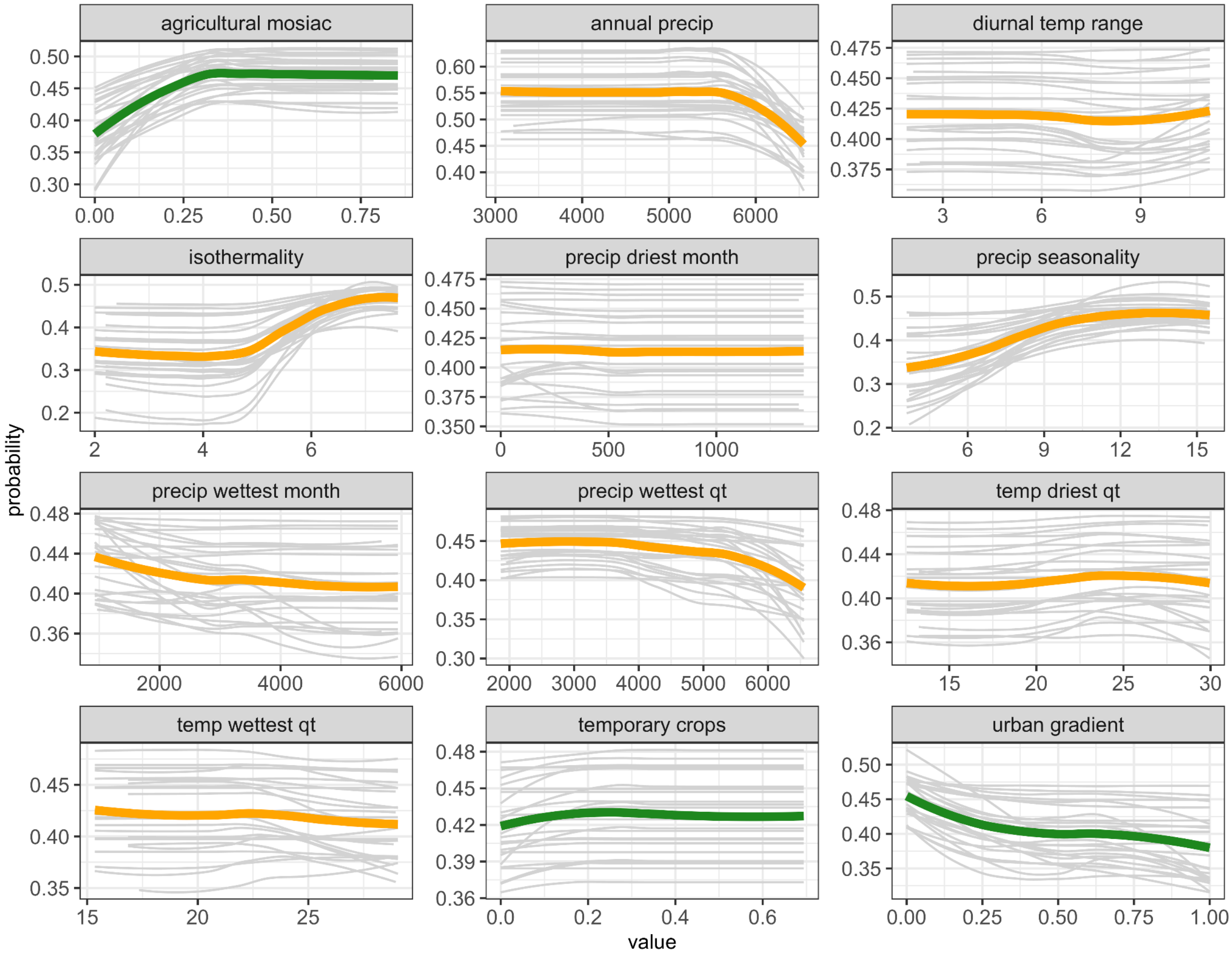


**Supplementary figure 13. *B. straminea* partial dependence plots for climate and land-use when the urban area is defined based on the UN definition.** Gray lines represent PDPs of each bootstrapping iteration, color lines represent average across iterations (orange = climate feature, green = land-use feature). Plots where the line looks flat indicate features with minimal contribution to feature importance.

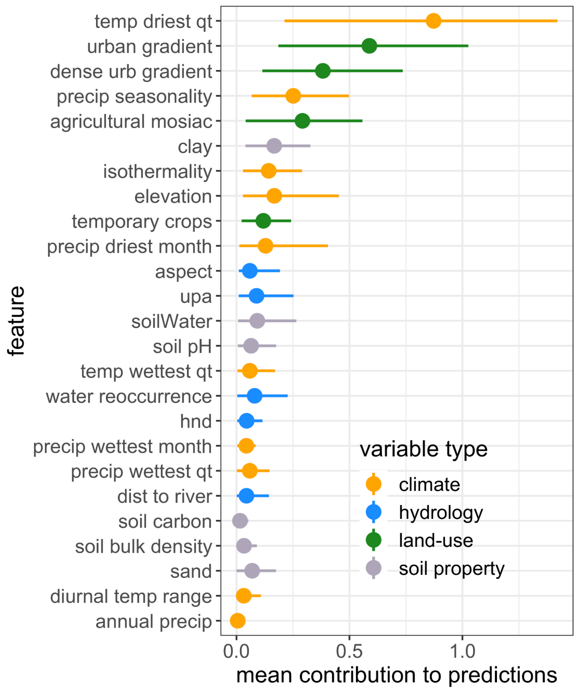


**Supplementary figure 14. Contributions to predictions for the *B. tenagophila* model when defining an urban area based on the UN definition.** Points are the absolute value of the mean contribution to predictions (mean |SHAP Value|) for the covariate across all data points (i.e., global feature contribution), bars represent the 95% confidence interval.

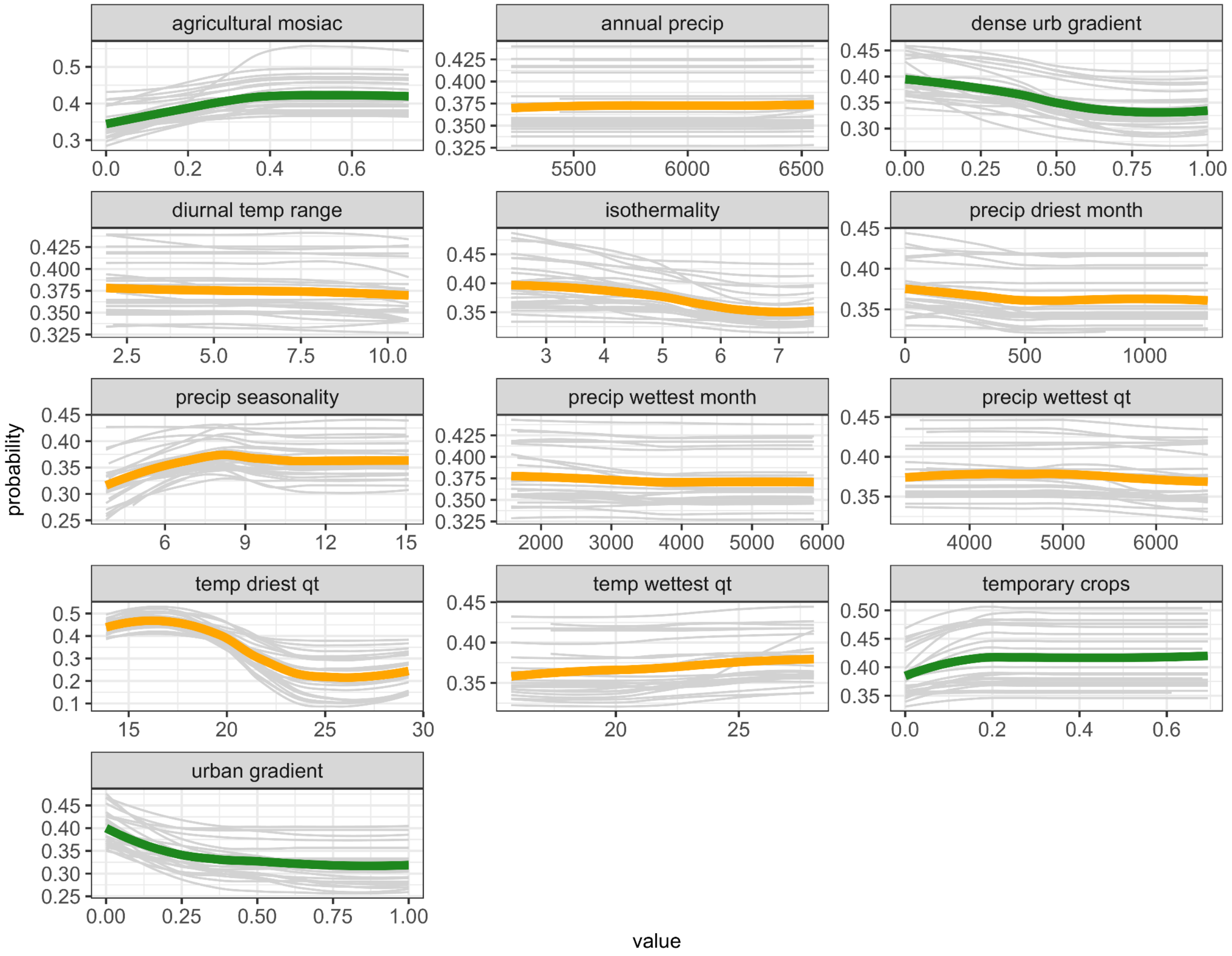


**Supplementary figure 15. *B. tenagophila* partial dependence plots for climate and land-use when the urban area is defined based on the UN definition.** Gray lines represent PDPs of each bootstrapping iteration, color lines represent average across iterations (orange = climate feature, green = land-use feature). Plots where the line looks flat indicate features with minimal contribution to feature importance.

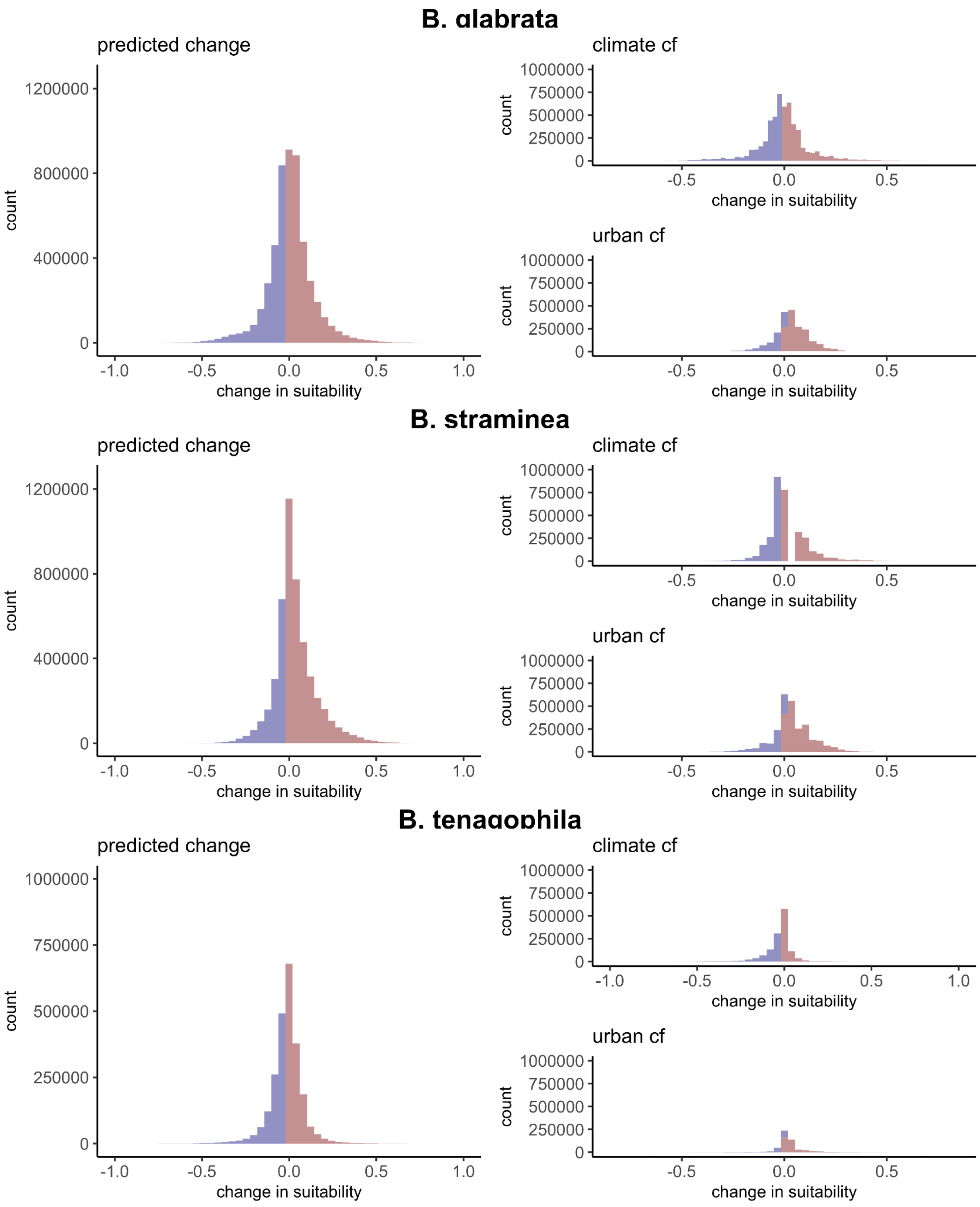


**Supplementary figure 17.** Total change in pixels associated with the change in suitability between 1992 and 2017, the suitability change in relation to the climate counterfactuals (2017 predictions - climate counterfactual predictions), and the suitability change in relation to the urban counterfactuals (2017 predictions - climate counterfactual predictions).

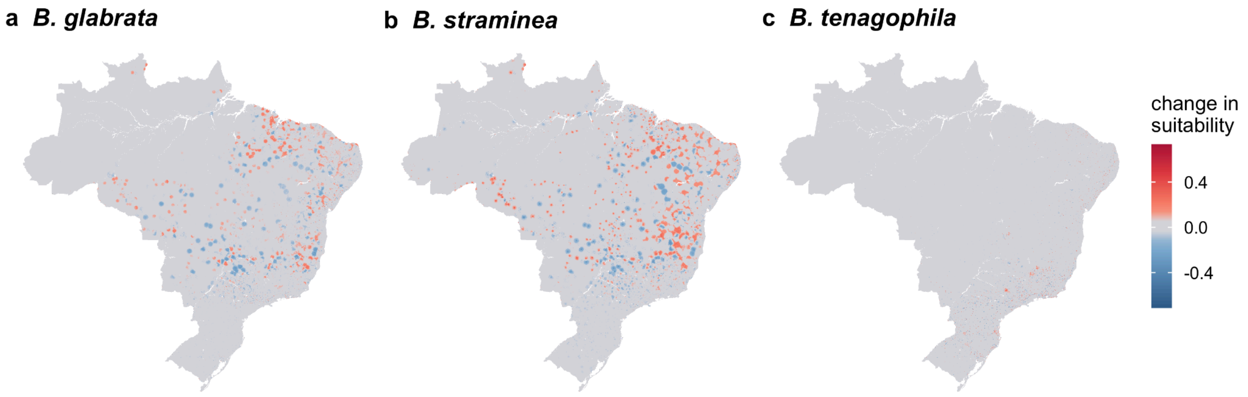


**Supplementary figure 18. Urban counterfactual analysis.** The difference in distribution between the predicted distribution of 2017 and when holding urban features constant (i.e., as they were observed in 1992). Blue indicates the habitat that became more suitable with urbanization and red indicates habitat became less suitable with the observed urbanization (i.e., the change in distribution is correlated with urbanization). Figures are mean values per pixel across 25 bootstrapping iterations.

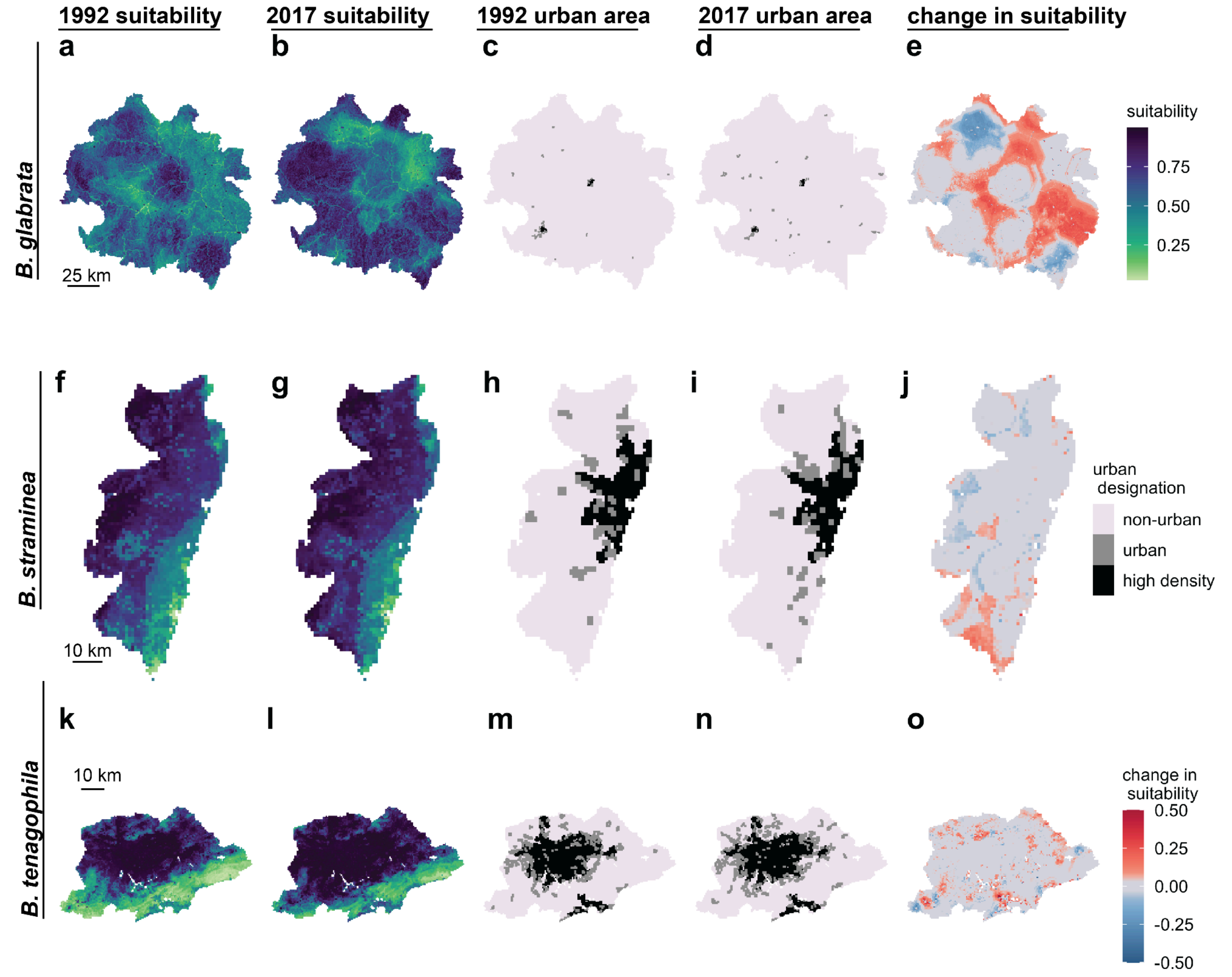


**Figure 19. Urbanization influences change in habitat suitability on small scales.** Predicted habitat suitability for each snail in 1992 **(a, f, k)** and 2017 **(b, g, k),** the urban extent of each mesoregion in 1992 **(c, h, m)**, and 2017 **(d, i, n)**, and change in suitability associated with urbanization **(e, j, o).** The *B. glabrata* maps are in the Vale de Rio Doce, the *B. straminea* maps are in the Recife Metropolitan Area, and the *B. tenagophila* maps are in the São Paulo Metropolitan area. High density = > 1500 people per 1km$^2$, with > 150,000 people in contiguous pixels; urban = > 300 people per 1km$^2$ pixel, with >2500 people in contiguous pixels.

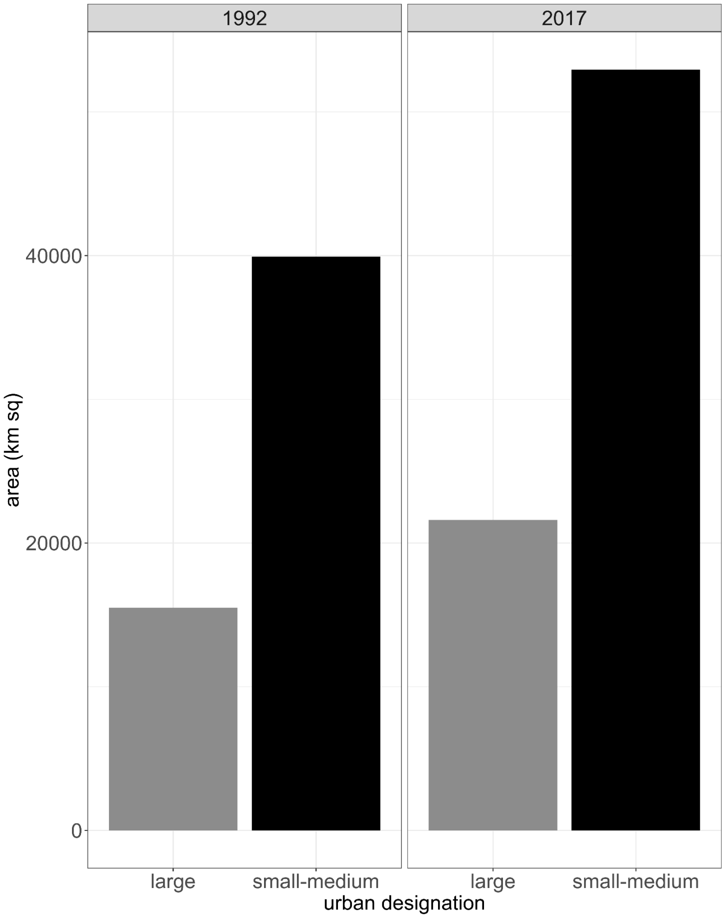


**Figure 20. Urban area ($km^2$) in 1992 and 2017.**
